# Coxsackievirus B3 Infection Early in Pregnancy Induces Congenital Heart Defects Through Suppression of Fetal Cardiomyocyte Proliferation

**DOI:** 10.1101/2020.05.19.104844

**Authors:** Vipul Sharma, Lisa S. Goessling, Anoop K. Brar, Chetanchandra S. Joshi, Indira U. Mysorekar, Pirooz Eghtesady

## Abstract

**Aims:** Coxsackievirus B (CVB), the most common cause of viral myocarditis, targets cardiomyocytes through Coxsackie and Adenovirus Receptor, which is highly expressed in the fetal heart. We hypothesized CVB3, a well-recognized culprit for viral myocarditis, can precipitate congenital heart defects (CHD), when fetal infection occurs during critical window of gestation.

**Methods & Results:** We infected C57Bl/6 pregnant mice with CVB3 during serial time points in early gestation (E5, E7, E9 and E11). We used different viral titers to examine possible dose-response relationship and assessed viral loads in various fetal organs as well as kinetics of virus passage into the fetus during gestation. Provided viral exposure occurred between E7-E9, we observed characteristic features of ventricular septal defect (33.6%), abnormal myocardial architecture resembling non-compaction (23.5%), and double outlet right ventricle (4.4%) among 209 viable fetuses examined. We observed a direct relationship between viral titers, severity and incidence of CHD, with apparent predominance among female fetuses. Infected dams remained healthy; we did not observe any maternal heart or placental injury suggestive of direct viral effects on developing heart as likely cause of CHD. We examined signaling pathways in CVB3-exposed hearts using RNAseq, KEGG enrichment analysis and immunohistochemistry. Signaling proteins of the Hippo, tight junction, transforming growth factor β1 and extracellular matrix proteins were the most highly enriched in CVB3-infected fetuses with VSD (log fold change >1.9, P<0.02). Moreover, cardiomyocyte proliferation was 50% lower in fetuses with VSD compared with uninfected controls.

**Conclusion:** Prenatal CVB3 infection can induce CHD, provided the infection occurs during a critical window. Alterations in myocardial proliferate capacity and consequent changes in cardiac architecture and trabeculation appear to account for the majority of observed phenotypes.

**GRAPHICAL ABSTRACT:** 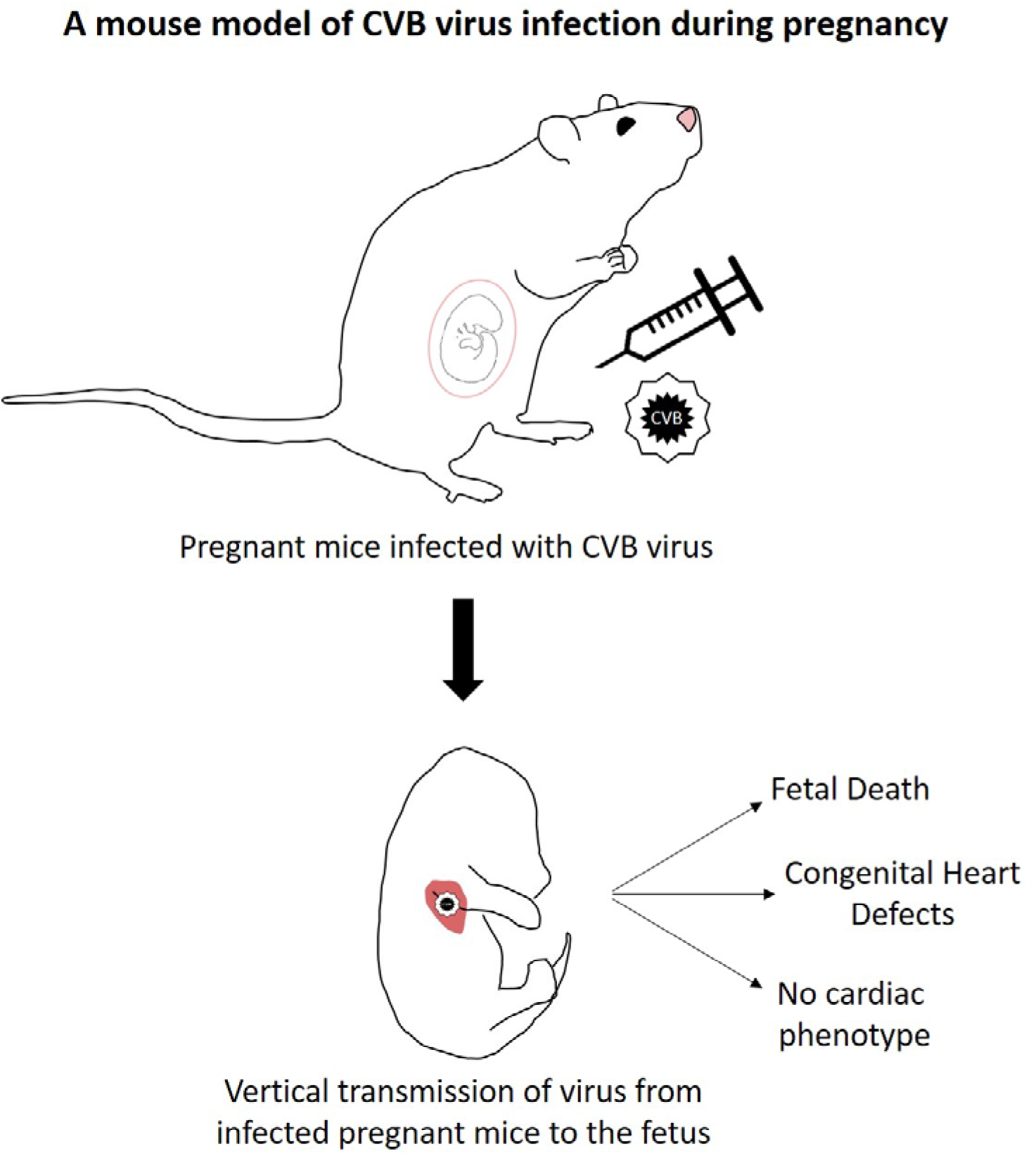

## INTRODUCTION

Congenital heart defects (CHD) are the most common anomalies in newborns, affecting 1 in 100 newborns annually ^[1]^. While many lesions are now treatable with excellent outcomes, still many suffer from a shortened life-span, while others endure lifelong residual cardiovascular problems ^[2]^, many of whom continue to need invasive treatments and not an insignificant number eventually requiring heart transplantation ^[3]^. Pedigree analysis of families with multiple affected individuals as well as other detailed genetic studies have provided insights into the genetic basis of several CHD, and potential heterogeneous pathogenic causes ^[4]^; however, for a significant portion of CHD the etiology remains largely elusive.

Since the Rubella epidemic of 1964 in United States, it has been recognized that viruses could play a role in etiology of many birth defects including CHD ^[5, 6]^. Some of these studies suggest a strong association between viral infections and fetal congenital anomalies of the nervous and cardiovascular systems ^[7]^. Overall, prenatal infections are presumed to account for 2-3% of all congenital anomalies, most often by infections with “TORCH” organisms: toxoplasmosis, rubella, cytomegalovirus, herpes and “other”^[8]^. The latter category of “other” pathogens is continually expanding: for example, a high incidence of congenital defects has been reported in infants born coincident with seasonal influenza epidemics and Zika virus infection during early pregnancy ^[9, 10]^.

Coxsackievirus group B (CVB) infects 10 million US citizens annually, allowing for a potentially large exposure burden. Six serotypes (B1–B6) are recognized, with serotypes B2, B3, and B4 considered endemic in the United States, and serotypes B1 and B5 exhibiting epidemic patterns ^[11]^. Coxsackievirus and Adenovirus Receptor (CAR, encoded by *CXADR* in humans) is believed to be the primary mediator of CVB infection, serving as a point for attachment and internalization for all CVB serotypes ^[12]^. *CXADR* is expressed in the human placenta during the first trimester ^[13]^ and transplacental transfer of CVB is well documented in humans ^[14]^, suggesting the potential for exposure of the developing fetus to viral infections. Evidence of CVB infection in late gestation mouse placenta and fetus is seen within 2-3 days of maternal infection, with a short duration of fetal viremia (2-4 days) ^[15]^. That the developing heart can be a target for virus-induced pathology is also suggested from studies showing that following perinatal maternal infection with CVB, viral RNA is detected in the neonatal heart for up to 5 days ^[16]^ and is associated with damage in neonatal cardiomyocytes ^[17]^. Studying serologic responses, Brown and Evans (1967) were the first to suggest a possible clinical association of early gestation maternal CVB infection with various congenital heart and brain defects, later posited by others as well ^[18-21]^. No further studies, however, have provided direct evidence that CVB causes CHD or a plausible mechanism to account for the clinical observations.

We have hypothesized that infection by “cardiotropic” viruses (i.e., viruses, such as CVB, that enter the cardiomyocyte via specific receptors in the fetal heart) can induce CHD by interfering with normal cardiomyocyte proliferation provided infection occurs during a critical window (first trimester) of cardiogenesis. To test our hypothesis, we used a mouse model to examine fetal exposure to CVB in early gestation. We also examined the influence of sex in this mechanism since prior studies have suggested sex differences in susceptibility to infection by CVB, as well as in the incidence of certain CHD ^[22, 23]^.

## MATERIAL AND METHODS

### Propagation of coxsackievirus and infectivity assays

LLC-MK2 Derivative cells (American Type Culture Collection [ATCC] CCL-7.1) were maintained in Minimum Essential Medium (MEM; Corning-Mediatech, 10-009-CV) containing 10% fetal bovine serum (FBS; Millipore, TMS-013-B) and 2mM (1X) GlutaMAX-I (Gibco, 35050-061) at 37°C in a 5% CO_2_ incubator. CVB3, Nancy (ATCC, VR-30) was propagated in monolayers of LLC-MK2, derivative cells. When CPE reached 95-100%, virus stocks were prepared by subjecting cells and media to three freeze-thaw cycles, clarifying the media by centrifugation (2800g, 10 minutes) and aliquoting the supernatant. Aliquots were quick frozen on dry ice and stored at -80°C. Virus concentration was determined using an endpoint dilution assay on LLC-MK2, derivative cells and expressed as a 50% tissue culture infective dose per milliliter (TCID50/ml) as calculated by the Spearman & Kärber algorithm ^[24]^. To determine the presence of infectious virus, fetal tissues were homogenized in serum-free Minimal Essential Media containing penicillin, streptomycin and amphotericin B. The suspension was frozen on dry ice, quickly thawed at 37°C then clarified by centrifugation at 2000xg for 10 min. A portion of the supernatant was used to inoculate LLC-MK2 derivative cells seeded the day prior in 24 well plates. Cultures were observed daily for the presence of CPE for at least one week. Live cells remaining on the plates following the observation period were fixed with 4% PFA and stained with 0.1% Crystal Violet.

### Experimental Animals

C57Bl6/J mice were purchased from The Jackson Laboratory and allowed to acclimate for one week prior to any procedures. Animal protocols were approved by the Institutional Animal Care and Use Committee at Washington University School of Medicine (protocol number 20170070).

All procedures conformed to the guidelines from the NIH Guide for the Care and Use of Laboratory Animals. Matings were set up overnight and the following morning, considered embryonic day 1 (E1), the females were examined for the presence of a copulation plug. Weights were monitored throughout the gestation period and weight gain was used as an indicator of the progression of pregnancy and maternal health post-infection. Pregnant females were randomly assigned into either a control group (n=7) or groups inoculated by intra-peritoneal injection with the virus (n=42) at the selected gestational day. We used a matrix with four time points of CVB3 infection (E5, E7, E9, and E11) and three viral inoculation doses (1 × 10^6^ TCID50, 2.5 × 10^6^ TCID50, and 5 × 10^6^ TCID50). Dams were observed daily following inoculation for signs of illness (i.e, lethargy, rough coat, etc.). Feto-placental units from all groups were collected by laparotomy under general anesthesia at the desired gestational stage. Dams were euthanized by exsanguination and cardiectomy under inhaled 5% isoflurane anesthesia. Placental and fetal tissues were either flash frozen, embedded in OCT (Tissue-Tek, 4583) then frozen or fixed in 4% paraformaldehyde (PFA) and embedded in paraffin for immunohistochemical (IHC) analysis. The fetuses were collected at E15-17, as normal cardiac formation would be complete and hence, any abnormalities identified would not be residua of incomplete development.

### Histology and Immunohistochemistry

Transverse serial 7-µm sections of the mouse fetus and placenta were stained with hematoxylin and eosin to assess heart morphology. Maternal heart sections were stained with Masson’s Trichrome stain to detect cardiac injury. Sections for IHC were deparaffinized and rehydrated, then antigen retrieval was performed using the citrate-based solution (H-3300) and high temperature-pressure protocol of Vector Laboratories (Burlingame, CA). Samples were blocked for 1hr at room temperature with 10% FBS then incubated overnight at 4°C with primary antibodies PHH3 (Millipore 06-570) to identify mitotic cells, MF20 (Developmental Studies Hybridoma Bank, MF20) to identify myocytes, and Mouse Anti-Enterovirus Clone 5-D8/1 (Dako M7064) to identify CVB VP1 region. For immunofluorescence detection, sections were incubated with Alexa Fluor conjugated secondary antibodies (Invitrogen, goat anti-rabbit IgG #A11011 and goat anti-mouse IgG #A11001) for 2 hours at room temperature. Slides were mounted with Vectashield Mounting media with DAPI (Vector Labs, H-1200) and imaged using a Zeiss Fluorescent microscope (Axiovert with Apotome 2 apparatus). Apoptosis in the experimental groups was assessed in a blinded manner by TUNEL staining using the *In Situ* Cell Death Detection Kit, TMR red (Roche 12156792910) according to manufacturer’s instructions.

### Placental histology

Placental sections obtained from infected and uninfected groups were processed for histological analysis. Hematoxylin and Eosin staining was performed on sections as per standard protocol. Three to four placental sections of the decidua, junctional zone and labyrinth zone from each group were examined in a blinded fashion (C.J. and I.M.) for evidence of pathology.

### Fetal Heart Culture

Fetal hearts were collected at E11 from uninfected and CVB3 infected (E9) dams and cultured on Millicell EZ 8-well slides (Millipore Sigma PEZGS0816) in high-glucose DMEM (ATCC 30- 2002) supplemented with 10% Fetal Bovine Serum (Millipore TMS-013-B), 1% Penicillin Streptomycin (Gibco 15140-122), and 1% Amphotericin B (Corning 30-003-CF). Cells were grown for 3 days, to achieve an age comparable to E14 gestation, and fixed with PFA for IHC.

### Mouse fetal sex determination

Genomic DNA was extracted from frozen tissue using Tissue Direct PCR kit (Lamda Biotech D300-100), amplified using Taq Polymerase 2X PCR premix (Intact Genomics, 3249) and SX primers (SX_F, 5’-GATGATTTGAGTGGAAATGTGAGGTA-3’; SX_R, 5’- CTTATGTTTATAGGCATGCACCATGTA-3’) ^[25]^ and the PCR products analyzed on a 2% agarose gel. The number of males and females in uninfected fetuses were used to exclude the possibility of skewed analysis from uneven sex distribution among conceptuses and the probability of females having the defect over males by using the formula:

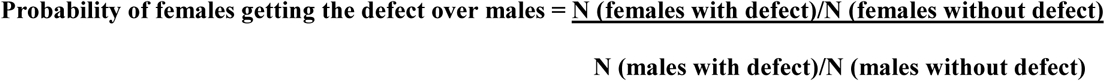

### Determination of cell proliferation

Cell proliferation in fetal hearts was determined by mitotic index, which was expressed as the percentage of pHH3 stained nuclei in MF20-positive cardiomyocytes/total number of nuclei in MF20-positive cardiomyocytes. Cell counting was calculated in 4 biological replicates using ImageJ image processing software (U.S National Institutes of Health) ^[26]^. Individual data points together with the average measurements are reported as the mean ± standard deviation (SD).

### PCR analysis and RNA-sequencing

RNA extraction was done using QIAamp Viral RNA mini kit (Qiagen 52904) for RT-PCR and Arcturus PicoPure RNA isolation kit (Applied Biosystems 12204-01) for qPCR and sequencing using manual guidelines. cDNA synthesis was done using SuperScript III First-Stand (Invitrogen 18080-051) and RT All-in-one master mix (Lamda Biotech G208-100). Endpoint PCR was done using EV primers (Forward – CGGCCCCTGAATGCGGCTAATCC, Reverse – TTGTCACCATAAGCAGCCA) ^[27]^. PowerUp SYBR Green Mastermix (Applied Biosystems A25741) was used for qPCR to quantify CVB (Forward – CCCCGGACTGAGTATCAATA, Reverse – GCAGTTAGGATTAGCCGCAT) ^[28]^. TGFβ1 (Forward CAACAATTCCTGGCGTTACCTTGG, Reverse – GAAAGCCCTGTATTCCGTCTCCTT), GAPDH (Forward GCATGGCCTTCCGTGTTC, Reverse – GATGTCATCATACTTGGCAGGTTT) ^[29]^, Bone morphogenetic protein 2 (BMP2) (Forward TGTGGGCCCTCATAAAGAAGCAGA, Reverse – AGATCCCTGCTTCTCAAAGGCACT) ^[30]^, Smad1 (Forward CGCTCCACGGCACAGTTAAG, Reverse – GCCAGTTGATTTGCGAACAGAA), Smad5 (Forward TGCAGCTTGACCGTCCTTACC, Reverse – GCAGACCTACAGTGCAGCCATC), and Smad9 (Forward CGATCATTCCATGAAGCTGACAA, Reverse – TGGGCAAGCCAAACCGATA) ^[31]^. For sequence analysis fetal tissues embedded in OCT were sectioned at 10µm onto polyethylene napthalate (PEN) membrane slides (Leica, 11505158) and heart tissue was collected using a Leica LMD7000 Laser Microdissection System from CVB3-infected fetuses showing a VSD and uninfected control fetuses. RNA was extracted from the heart tissue and submitted to the Genome Technology Access Center (GTAC, Department of Genetics at Washington University in St. Louis) to perform RNA-Seq using next generation sequencing for transcriptome profiling. The Kyoto Encyclopedia of Genes and Genomes (KEGG) database was used to identify biological pathways linked to differentially expressed genes enriched in CVB3-infected fetal hearts.

### Statistical Methods

T-test was used for continuous values and comparisons of categorical variables were made by calculating median and by Chi-square test to analyze statistical significance between control and test groups by calculating p values (value less than or equal to 0.05 was considered statistically significant). For multiple biological replicates standard deviation was calculated. To quantify amount of virus in tissue samples standard curve equation was used along with its coefficient of determination (R^2^).

## RESULTS

### CVB3 passes from infected pregnant dam into the fetuses

To demonstrate passage of CVB3 from the infected mother to the developing fetus initially we performed infectivity assay and endpoint PCR on fetal lysates (Figure 1 a, b). To confirm the presence of virus within the fetus (and to localize the viral particles), we made multiple attempts at RNA in-situ hybridization without success. We therefore, used qPCR not only to confirm the presence of the virus but also to quantify the viral load within the fetus. Using a standard curve based on our CVB3 stock (Figure 1d), we measured CVB3 levels in placenta (Figure 1e) and various fetal organs (Figure 1f) at different time-points (E11, E14, and E17) following inoculation at E9. This data shows that high levels of virus persist for at least 6 days within the placenta. CVB3 quantification in different fetal organs was done with specific emphasis on those which express CAR. Since dissection of individual organs was not possible at E11, we elected to divide the fetus into head (brain), mid-torso (heart), and lower-torso (liver, intestine) segments for analysis at that gestational age. At E14, we examined heart, brain, mid-torso (predominantly liver), and lower- torso. At E17 we could dissect out all individual organs. We found the maximum amount of CVB3 in the fetal heart at E14 followed by fetal brain at E17 (Figure 1f).

**Figure 1.**
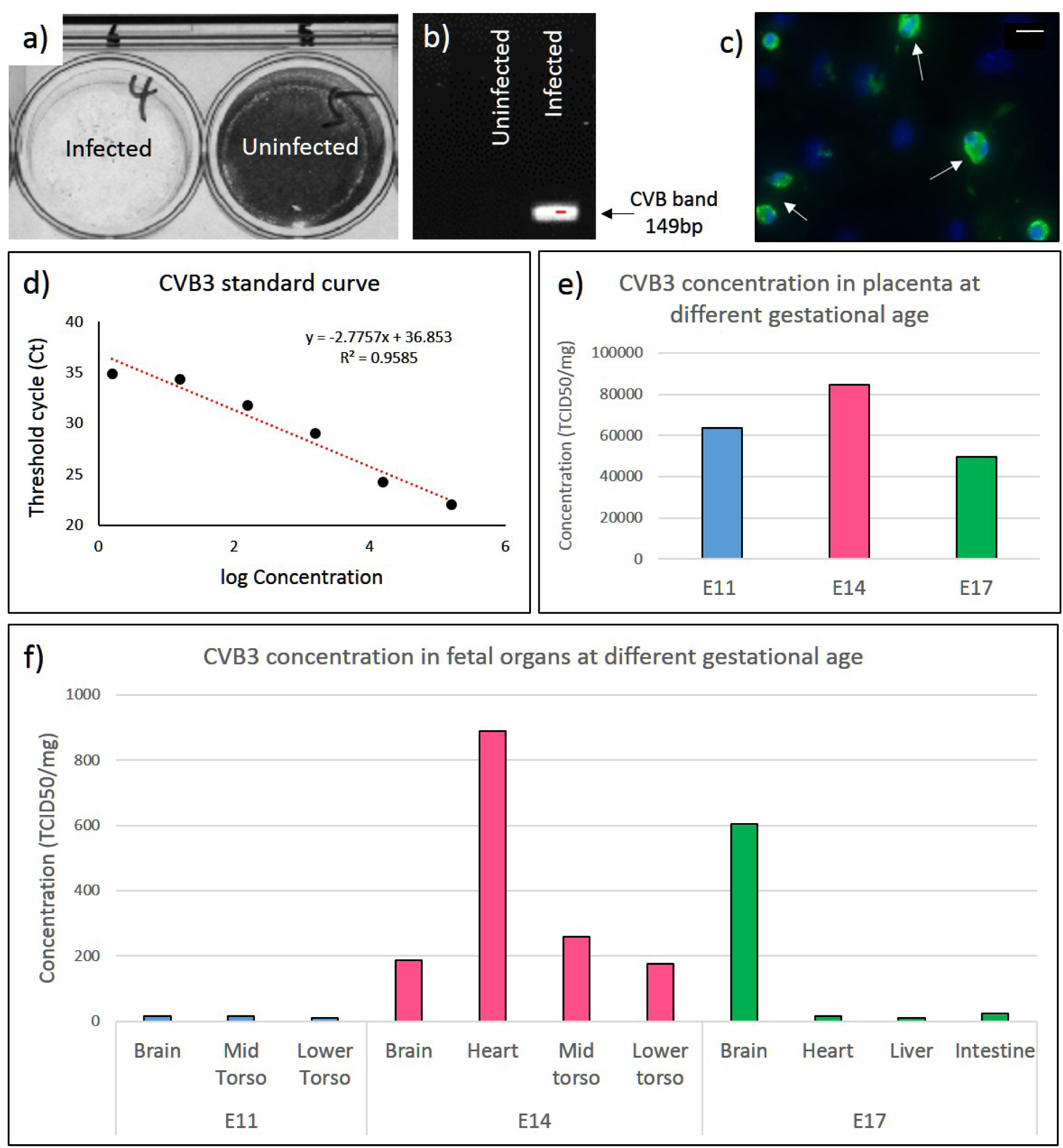
Vertical transmission of CVB3 from infected pregnant dams to the fetus. a) The presence of infectious virus in fetal lysates from infected and uninfected dams (E9) was analyzed using a cell culture infectivity assay; the lack of stained cells indicates the presence of virus. b) Reverse Transcriptase-PCR with enterovirus (EV)-specific primers showing amplification of the 149-bp (determined by agarose gel electrophoresis) virus band only in the tissue lysate of a fetus from an infected dam (at E9). c) IHC using anti-EV antibody on cultured fetal heart (∼E14) explanted at E11 from a dam infected at E9 (white arrows). Virus quantification by qPCR; d) Standard curve based on the concentration of CVB3 stock. e) virus present in placenta at E11, E14, and E17 after infection at E9. f) virus present in fetal organs at E11, E14, and E17 after infection at E9. To quantify the amount of virus in tissue samples, the standard curve equation was used along with its coefficient of determination (R^2^). Scale 20µm.

We also performed IHC to detect viral capsid protein (VP-1) on both fetal tissue sections and explanted hearts (at E11) that were cultured for 3 days (∼E14). While the anti-VP1 antibody did not reliably detect the virus in tissue sections of the fetal hearts, with in vitro culture we could easily detect VP-1, confirming the continued presence of the virus in the hearts of fetuses from CVB3 infected dams (Figure 1c).

### Maternal CVB3 infection leads to abnormal heart development in fetal mice

At E17, approximately one-third of the fetuses from CVB3 infected dams were either dead or resorbed (37.2%, n=124/333; p=0.001) and could not be processed for assessment of heart morphology (Figure 2a, f). Of the 209 infected fetuses assessed, 78 (37.3%; p=0.00002) had one or multiple CHD: 66 (31.6%; p=0.0005) exhibited ventricular septal defect (VSD) (Figure 2c, f; Supplementary Figure 1a-c for more representative sections), 48 (23%; p=0.00002) exhibited alteration in myocardial architecture grossly resembling non-compaction (NC) of the ventricular myocardium (Figure 2d, f; Supplementary Figure 1d-f for more representative sections), and 9 (4.3%; p=0.2) exhibited double outlet right ventricle (DORV) phenotype (Figure 2e, f). Of all the VSD seen, most were perimembranous in nature and only 6% were muscular VSDs (Supplementary Figure 1a). The viral loads determined in the fetuses showed resorbed fetuses had the highest viral loads per mg tissue, followed by fetuses with multiple CHD, and the lowest load seen in those with isolated VSD or NC (Figure 2 table). No other major heart pathology was noted.

**Figure 2.**
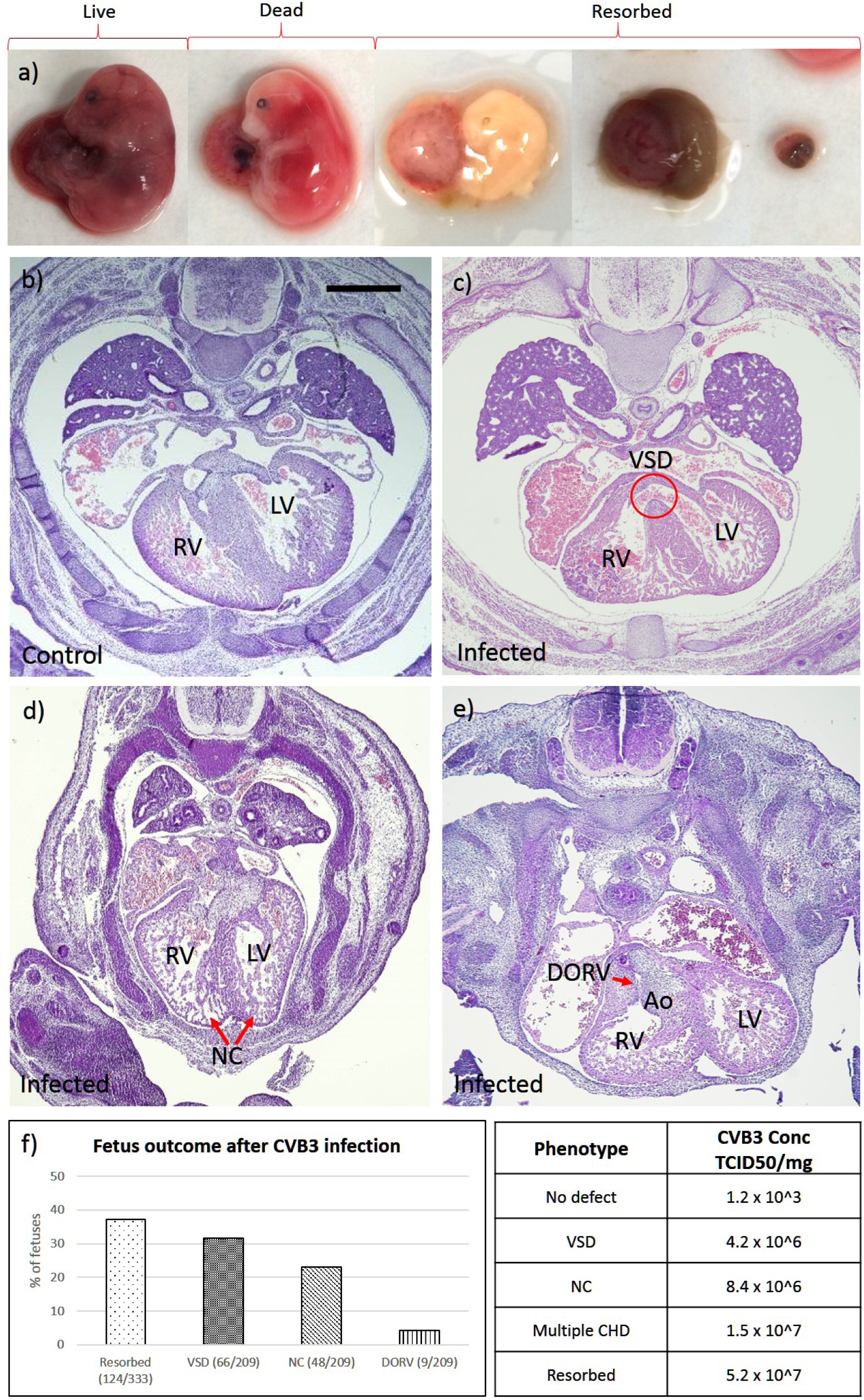
CVB3 infection of pregnant dams leads to fetal cardiac defects and death. a) Representative images of E17-fetuses from CVB3-infected dams exhibiting specific phenotypes (left to right): viable, fully formed dead, and 3 images showing fetuses at various stages of resorption. b–e) Hematoxylin and eosin-stained heart sections of E17-fetuses from uninfected (b) and infected dams (c, d, e), and the various cardiac defects are indicated. f) Graph showing the percentage of fetuses from CVB3 infected dams with the indicated cardiac defects. Chi-square test was used to analyze statistical significance between control and test groups by calculating p values. Table showing the amount of virus present in resorbed fetuses and fetuses with CHD determined by qPCR using CVB3 quantification standard curve (Figure 1e). RV = right ventricle; LV = left ventricle; Ao = Aorta; VSD = ventricular septum defect; DORV = double outflow right ventricle; NC = non-compaction. Scale 100µm.

### E7-9 is the critical window for CVB3 induction of CHD in fetal mice

Since expression of CAR, the receptor for CVB, is essential between E10–E11 for cardiomyocyte development ^[32]^, we determined the time in gestation when CVB infection had the greatest impact on the incidences of the induced CHD phenotype, based on the timeline of mouse cardiac development (Figure 3a). The highest proportion of fetuses exhibiting VSDs occurred after infection at E9, regardless of the viral dose, with even low doses of CVB3 infection leading to abnormal heart development (Figure 3b; p=0.00001). The same pattern was observed for fetuses with NC and DORV with more incidences observed after infection at E9 for all CVB3 doses (Supplementary Figure 2). Similarly, while fetal resorption occurred following CVB infection at each gestational stage, the most cases followed infection at E9 (Figure 3c; p=0.00001).

**Figure 3.**
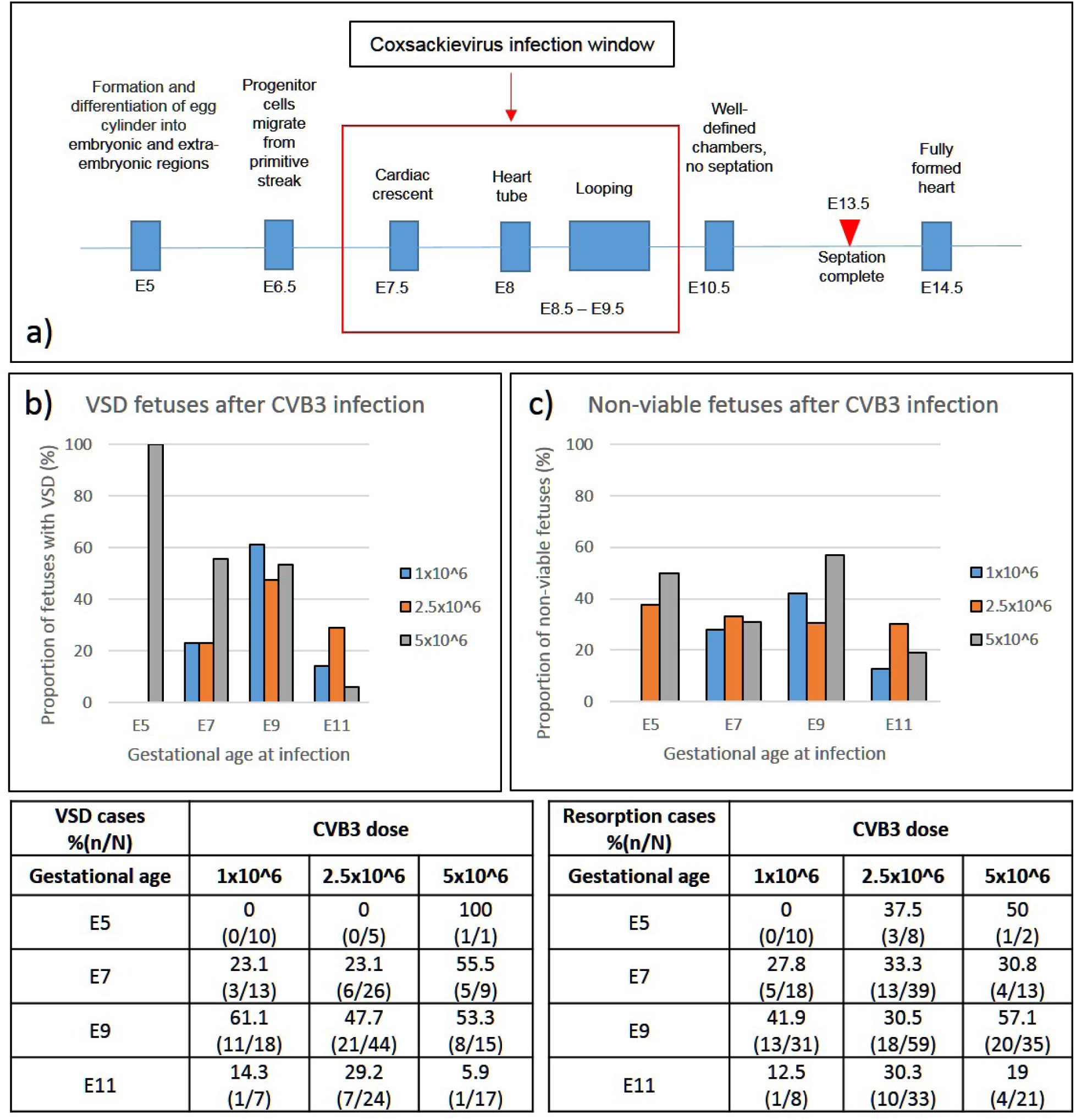
The effect of viral dose and gestational age at infection on the incidence of VSD and fetal demise. a) Schematic representation of the timeline of heart development in the mouse and critical window for CVB3 infection. b, c) Dams were infected at various stages of gestation (E5, E7, E9 or E11) with different doses of virus (1.0, 2.5 and 5.0×10^6^ TCID50). Graphs and their corresponding tables show the percentage of fetuses with VSDs (b), and non-viable (dead or resorbed) fetuses (c) along with the total examined in each experimental group. Chi-square test was used to analyze statistical significance between control and test groups by calculating p values.

### CHD is due to direct effect of CVB infection on the fetuses

To confirm the CHDs observed were a direct effect of CVB infection of the fetuses as opposed to poor health of the moms or placental pathology, we evaluated maternal health and examined placental histology. Infected dams did not show outward signs of illness such as lethargy or rough coat following CVB3 infection. We compared maternal weight gain pre- and post-infection between infected dams and uninfected control dams and found equivalent weight gain between the two groups (data not shown). Masson’s Trichrome staining of maternal heart tissue from control and infected dams also showed no evidence of myocarditis or cardiac injury (figure 4a-f). Placentas from control and E9 infected fetuses collected at E11 and E17 were analyzed by H&E staining and examined blindly by a placental biology expert (C.J. and I.M.). No placental damage, insufficiency, or vascular injury was identified (figure 4g-l). These results suggest the observed CHD are likely not simply a result of poor maternal health or placental insufficiency.

**Figure 4.**
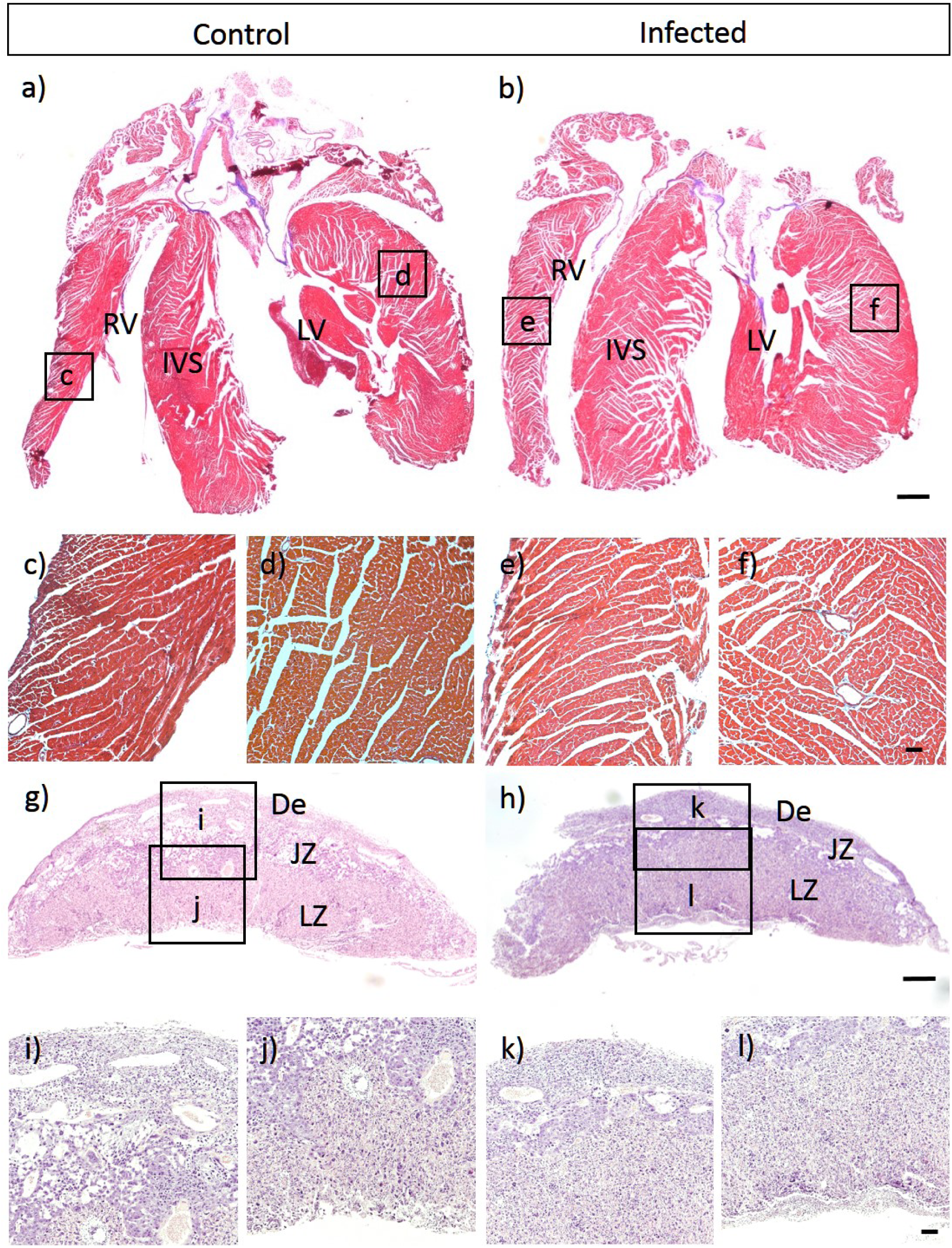
Cardiac defects in fetuses are not secondary to poor maternal health. a-f) Masson Trichrome stained heart sections of uninfected control (a) and infected dam (b). Higher magnification of control RV (c), control LV (d), infected RV (e), and infected LV (f) show no evidence of cardiac injury. g-l) Hematoxylin and eosin-stained placental sections of E17-fetuses from uninfected (g) and infected dams (h). Higher magnification of control (i,j) and infected (k,l) show no evidence of placental damage in different parts of placenta. RV = right ventricle; LV = left ventricle; IVS = inter ventricular septum; De = decidua; JZ = junctional zone; LZ = labyrinth zone. Scale 1000µm.

### CVB3 stimulates the TGFβ1 signaling pathway and suppresses cardiomyocyte proliferation

We found the maximum amount of CVB3 at E14 (Figure 1f); therefore, we used this time point for KEGG enrichment analysis of differentially expressed transcripts in VSD-exhibiting CVB3- infected and uninfected fetal hearts. Transcriptome analysis detected 13,118 protein-coding genes in the fetal heart, and differentially expressed genes were mapped to determine which were up- or down-regulated due to CVB3 infection (Supplementary Figure 3). The top five pathways most significantly enriched in CVB3-infected hearts (log fold-change >1.9, *P*<0.02) were those in the Hippo signaling, tight junction, TGFβ1, cell cycle, and extracellular matrix (ECM)-receptor pathways (Figure 5a). Number of genes up-regulated in these individual pathways were: 4 out of 118 genes in Hippo signaling, 3 out of 119 tight junction, 7 out of 61 TGFβ1, and 10 out of 61 in ECM receptor pathways, respectively.

**Figure 5.**
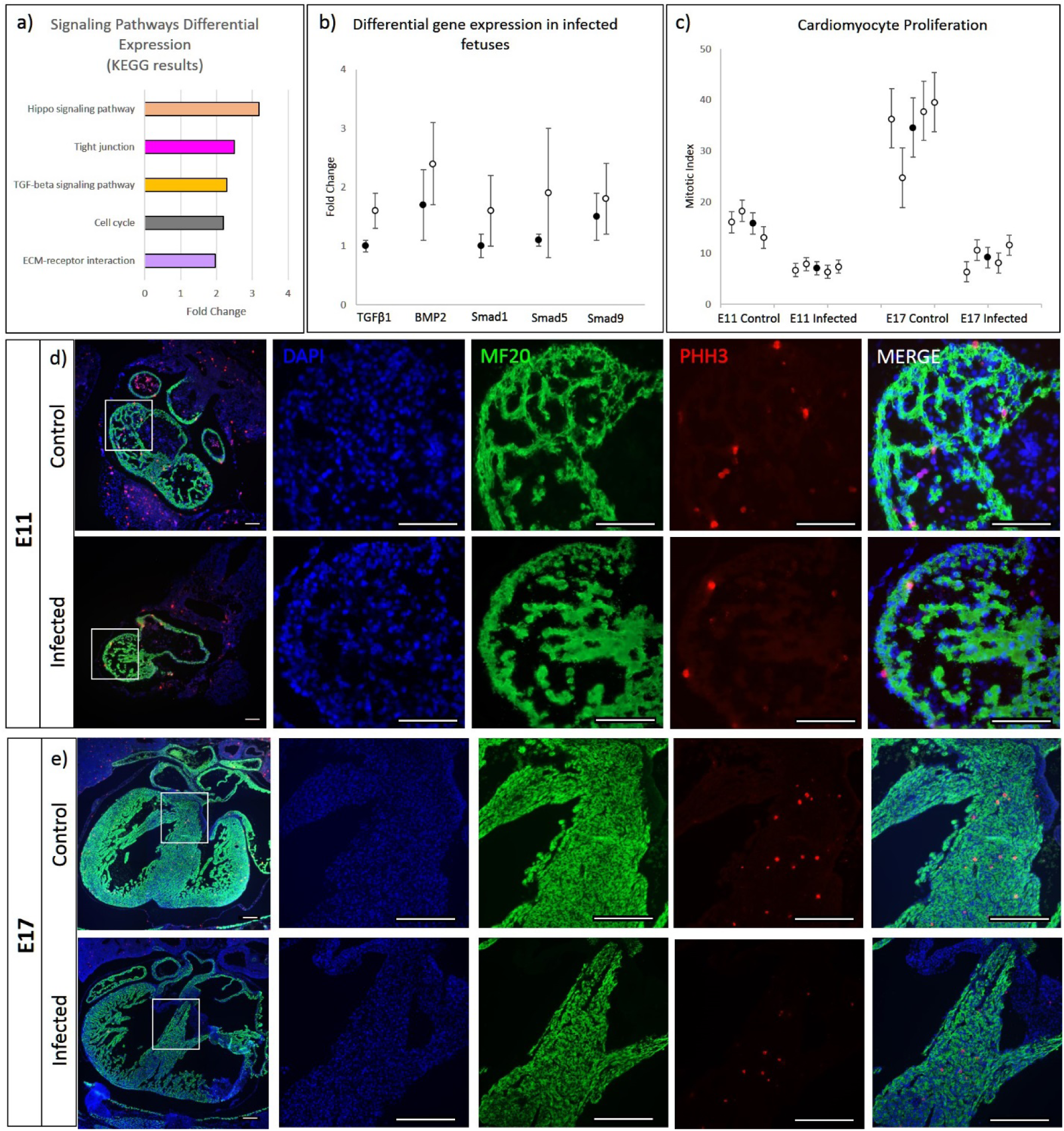
The effect of CVB3 infection on different signaling pathways and cell proliferation. a) Top five differentially regulated KEGG biological pathways in heart tissue from E14 fetuses infected with CVB3 at E9. b) Comparison of RNA sequencing (black dot, n=4) and RT-qPCR (white dot, n=4) data for up-regulated genes within the pathways (TGFβ1, BMP2, *Smad1, Smad5*, and *Smad9*), normalized to *Gapdh* expression in heart tissue from E14 fetuses infected with CVB3 at E9. c) pHH3-stained cells (individual data points as white dots and mean as black dot ± SD) indicating lower cell proliferation in the CVB3 infected fetal hearts (E11 and E17) compared to uninfected controls. d, e) Immunofluorescence images showing DAPI-stained DNA (blue), MF20-stained cardiomyocytes (green) and pHH3-stained proliferating cells (red) in the E11 control and infected fetuses (d) and E17 control and infected fetuses (e) (magnified images of the white box in the panels on the left). The number of stained cells was determined using ImageJ software. The images are representative of 4 biological replicates. Scale 100µm.

Since increased TGFβ1 expression has been reported in a number of CVB3 infection related models ^[33]^, and because the expression of TGFβ1 transcript has been found in regions actively undergoing cardiac septation in murine embryos ^[34]^, we examined whether the expression of other members of the TGFβ superfamily known to be associated with cardiogenesis were altered. Quantitative PCR was used to validate differentially expressed genes and the findings were consistent with data obtained by RNA sequencing. In particular, we found upregulation of genes and transcripts encoding TGFβ1, BMP2, and its downstream mediators SMAD1/5/9 in fetal hearts with VSDs (Figure 5b).

Based on the preceding findings, we assessed cell proliferation and apoptosis at E11, focusing on a developmental time point before septal completion (E13.5) (Figure 3a). We also examined E17 fetal heart sections to determine if the effects of CVB3 infection on proliferation were temporary or persistent. In heart sections stained with an antibody against phosphorylated histone H3, a marker for mitosis, cell proliferation in fetuses infected at E9 and collected at E11 and E17 was significantly lower than that in control fetuses with a mitotic index of 7.8% vs 15.8% at E11 and 9.1% vs 36.6% at E17 (p=0.0006 and 0.0002 respectively) (Figure 5c, d, e). Cell apoptosis levels assessed by TUNEL staining were similar in CVB3-infected and control fetuses (data not shown).

### Female mouse fetuses appear more susceptible to CVB3-induced CHD than males

Since sex differences have been reported in the prevalence of congenital anomalies and specific cardiac defects ^[35]^, we examined whether there were gender differences with CVB3-induced CHD. We found more female fetuses developing CHD than males, though not reaching statistical significance (Figure 6). A 1.5-fold higher (p=0.1) VSD incidence among females was observed compared with males. Females appeared also more susceptible to DORV and NC after CVB3 infection than males, with a 3.4-fold (p=0.1) and 1.4-fold (p= 0.2) increase respectively. Further, 1.2-fold (p=0.4) more female fetuses were resorbed compared to males following in utero CVB3 infection, suggestive of perhaps greater susceptibility of the female fetuses to infection.

**Figure 6.**
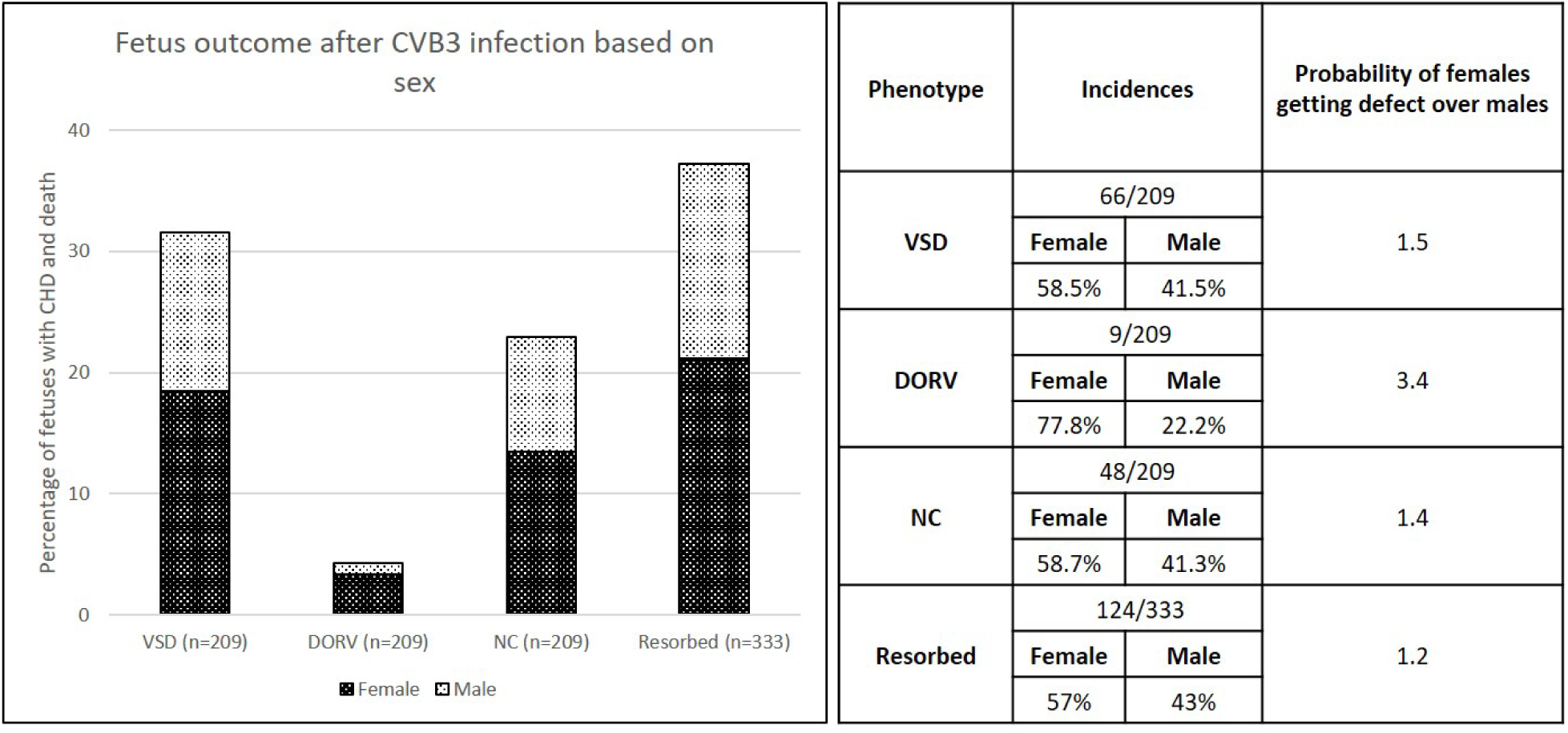
Difference in incidence of observed defects following CVB3 infection based on sex. The percentage of fetuses from CVB3-infected dams exhibiting the indicated abnormal phenotype. Chi-square test was used to analyze statistical significance between control and test groups by calculating p values.

## DISCUSSION

Viral infections in early pregnancy are recognized as important factors that can affect cardiovascular development ever since an association between intrauterine rubella infection and CHD was first reported in humans over 45 years ago ^[36]^. Similarly, suggestive, although inconclusive, evidence had been reported associating coxsackievirus infection during pregnancy and CHD ^[37]^. In the current study, we show conclusively prenatal CVB3 infection can indeed induce CHD, confirming prior suspicions based on epidemiologic studies. Moreover, we find viral load to directly relate to both severity and incidence of fetal pathology, showing highest viral loads in the resorbed fetuses, followed by fetuses with multiple CHD, and lowest virus levels in fetuses with isolated CHD. The major heart defect observed in our study, occurring in ∼30% of fetal mice exposed to CVB *in utero*, was perimembranous VSD, one of the most prevalent CHD in humans. Though not the focus of our study, it is of interest the second fetal organ where we identified abundant levels of CVB3 was the brain. Some studies have suggested the co-occurrence of central nervous system pathologies with CHD, though often these are attributed to secondary effects of altered in utero circulation from the primary CHD. Lastly, we identified a critical window during mouse development (between E7-9) when prenatal CVB3 infection has the greatest impact on inducing heart defects.

Prior reports using late gestation fetal mice suggested concentrations of CVB3 in fetal tissues peak at approximately 3 days following maternal infection ^[38]^. We observed a similar pattern therefore speculate maternal CVB3 infections between E7–9 in our experiments resulted in peak viral concentrations in the fetal heart around E10-12, when septation of the primitive cardiac tube is completing. Exposure of the fetal heart shortly before or during the period of septation could explain the temporal and quantitative relationship between CVB3 infection and the incidence of VSD that we observed in fetal mice. Since we did not observe any obvious placental injury and because the infected moms remained healthy throughout pregnancy, we suspect the heart defects resulted directly from infection of the fetus rather than secondary to placental insufficiency or poor maternal health. Our viral quantification studies suggest the placenta does indeed perform a superb job at filtering much of the viral wave. Some viruses, however, clearly manage to get across the placenta and persist for some time in the fetus with preferential proliferation in certain tissues (e.g., heart and brain).

It is of interest we also observed abnormal trabeculations and myocardial architecture resembling NC in CVB3 infected fetal hearts. Non-compaction of the left ventricle (LVNC), diagnosed in some children with CHD ^[39]^, can be isolated or associated with other congenital cardiac malformations. Although the etiology of NC is presently unknown, LVNC is attributed to arise from the arrest of the normal myocardial maturation process during ontogenesis, with familial or genetic triggers ^[40]^. Despite some controversy regarding diagnosis of NV, most agree NC is a genetically and phenotypically heterogenous disease that can affect both ventricles, and has been associated with several genetic variants as well as mutations in cytoskeletal and sarcomeric proteins ^[41]^. While our data showing that approximately 25% of CVB3-infected fetal hearts develop abnormalities of trabeculation somewhat similar to NC, perhaps due to CVB3 mediated arrested myocardial development, additional studies are required to correlate our findings to those observed in the clinical setting as well as potential mechanisms involved.

We examined signaling pathways to obtain insights into the molecular mechanisms responsible for the observed phenotypes in our mice, focusing on differentially regulated genes as a result of CVB3 infection. Enrichment of genes encoding for components of the TGFβ1 signaling pathway in fetuses with CVB3-induced heart defects, like elevated expression of BMP2 and its downstream signaling transcription factors, Smad 1, 5, and 9, may be related to observed VSD as well as altered trabeculation patterns in our mice. Increase in TGFβ1 expression is associated with cardiomyopathies in general but also with CVB3 induced dilated cardiomyopathy ^[33, 42]^. A critical role for BMP signaling in cardiac development has been extensively studied; deletion of *Bmp2* from the endocardial lineage results in perimembranous VSDs in mouse embryos ^[43]^. Moreover, up-regulation of BMP2 in the cardiac fields phosphorylates SMAD1/5/9 and enhanced SMAD signaling suppresses cell proliferation ^[44]^. We found CVB3 suppressed cardiomyocyte proliferation in the fetal heart suggesting it is one of the possible reasons behind observed phenotypes in our mice ^[45]^, and the resultant septation (VSD) and alignment defects (DORV) (Figure 7).

**Figure 7.**
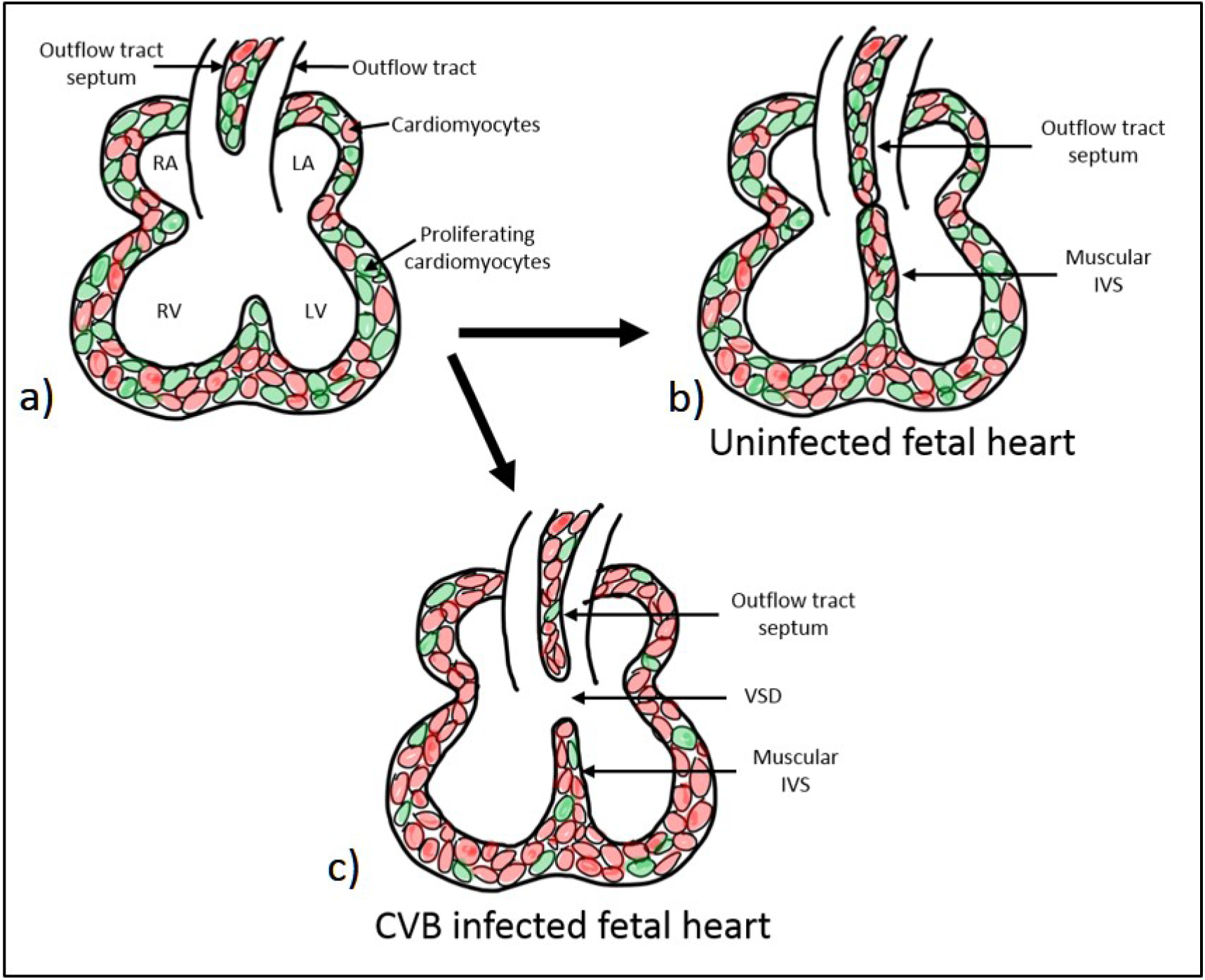
Proposed mechanism for VSD formation in the mouse model after CVB3 infection. a–c) The proposed mechanism of VSD generation. Normal intraventricular septum formation (a), with the proliferation of the muscular intra-ventricular septum and the outflow tract septum resulting in a fully formed septum (b), are shown (proliferating cardiomyocytes are shown in green). c) In CVB-infected fetuses, cell proliferation is suppressed, which leads to incomplete septum formation and VSD.

KEGG pathway enrichment analysis also showed that genes in the Hippo signaling pathways, known to regulate cardiovascular development and function, were enriched in CVB3-infected fetal hearts compared with uninfected controls. The Hippo signaling pathway is conserved across species and has a critical role in regulating embryonic organ (including heart) size and cardiomyocyte cell proliferation and apoptosis ^[46]^. Our observations of altered fetal cardiomyocyte proliferate capacity is therefore consistent with up-regulation of Hippo in our model. We also found expression of tight junction signaling pathways to be enriched in fetal hearts exposed to CVB3. Of note, the receptor for CVB3, CAR, is highly expressed in the developing murine heart and CAR knockouts result in embryonic lethality from cardiac defects if expression is repressed during a critical developmental window (E10 and E12) ^[32]^. This critical window coincides with the time at which fetal hearts in our model would be exposed to the highest levels of CVB3 following maternal infection at E7-E9. Although our transcriptomic data showed equal expression of CAR and ZO-1 in CVB3 infected and uninfected heart tissue (data not shown), we plan to pursue further analysis of this pathway components. With regard to all transcriptomic studies it is critical to changes at the protein level may still occur without seeing a change in transcript levels. In totality, however, our RNAseq data does suggest CHD observed in our model is likely related to alterations in proliferation capacity of fetal cardiomyocytes from CVB3 infection.

Sex hormones are thought to affect the susceptibility or response to CVB3 infection as well as CVB3-induced myocarditis in adult mice. In the preceding studies, males show more susceptibility to effects of infection than females ^[47]^. Moreover, testosterone has been shown to alter cardiac remodeling during CVB3 induced myocarditis ^[48]^. We observed some semblance of sex- influenced differences as well, albeit the reverse of preceding observations; however, despite the rather large number of fetuses examined, our observed ratios did not reach statistical significance. Clinicians and epidemiologists have reported unequal sex distribution of various cardiac defects in clinical cohorts ^[23, 35]^ though frequently with conflicting results: equal male/female ratios for VSD, but higher DORV incidence in males than females ^[49]^ versus results from a large Dutch study of 4110 adults with CHD, the incidence of VSD was higher among females (53%) than males ^[50]^. Female fetuses are often over-represented in stillbirth and recurrent miscarriage cases ^[51]^. We are interested to further dissect the influence of gender in our model in future studies as likely these subtle differences reflect as of yet not understood but important mechanisms.

In conclusion, while birth malformations caused by maternal viral infections, such as congenital rubella syndrome and those by the Zika virus have been well documented ^[10]^, we present novel data that maternal infection with a common enterovirus (CVB) can also result in development of structural heart defects. We found differential regulation between the uninfected and CVB-infected fetal hearts of signaling pathways associated with cardiac development. Based on these findings, we posit BMP/SMAD2 signaling-led suppressed cardiomyocyte proliferation as one possible mechanism of CHD pathogenesis associated with CVB infection. We believe our findings could have simple but important implications from the perspective of possible public health measures to reduce the incidence of CHD if clinical observations corroborate our findings.

## ACKNOWLEDGEMENTS

We thank Connor Mullen, Alma Muller, Brian Dailey, and Daniel Perry for data collection and histology support.

## AUTHOR CONTRIBUTION

PE devised the project and the main conceptual ideas, and supervised the project.

VS, LSG, AKB, PE conceived and planned the experiments.

VS and LSG performed the experiments.

VS analyzed the data and wrote the manuscript (with input and feedback from PE, AKB, LSG, IUM).

CSJ and IUM performed placental histopathological analysis.

## SUPPLEMENTARY DATA

**Supplementary Figure 1.**
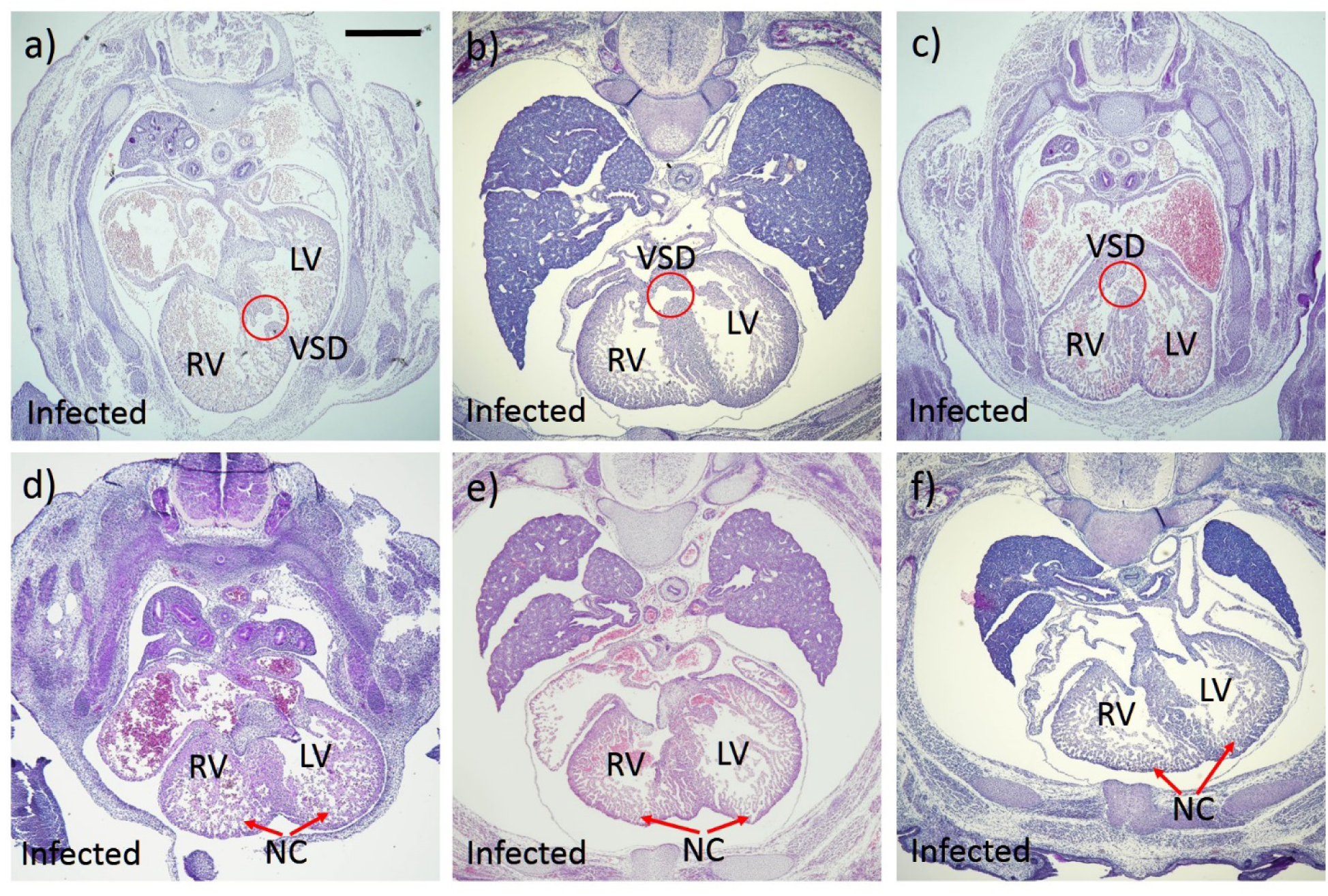
CVB3 infection of pregnant dams leads to VSD and NC. Hematoxylin and eosin-stained heart sections of E17-fetuses from infected dams with indicated cardiac defects: muscular VSD (a), perimembranous VSD (b, c), NC (d, e, f). RV = right ventricle; LV = left ventricle; VSD = ventricular septum defect; NC = non-compaction. Scale 100µm.

**Supplementary Figure 2.**
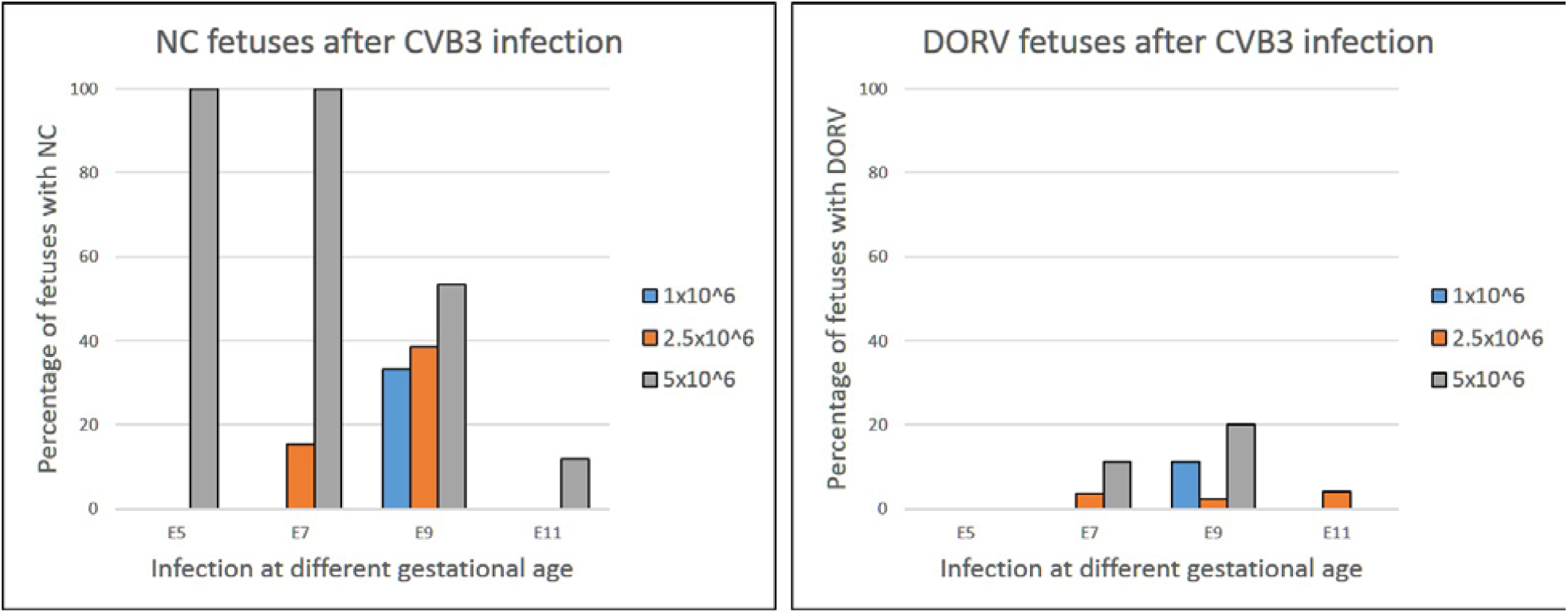
The effect of viral dose and gestational age on infection on the incidence of NC and DORV. Dams were infected at various stages of gestation (E5, E7, E9 or E11) with different doses of virus (1.0, 2.5 and 5.0×10^6^ TCID50). Graphs show the percentage of fetuses with NC (a) and DORV (b) of the total examined in each experimental group.

**Supplementary Figure 3.**
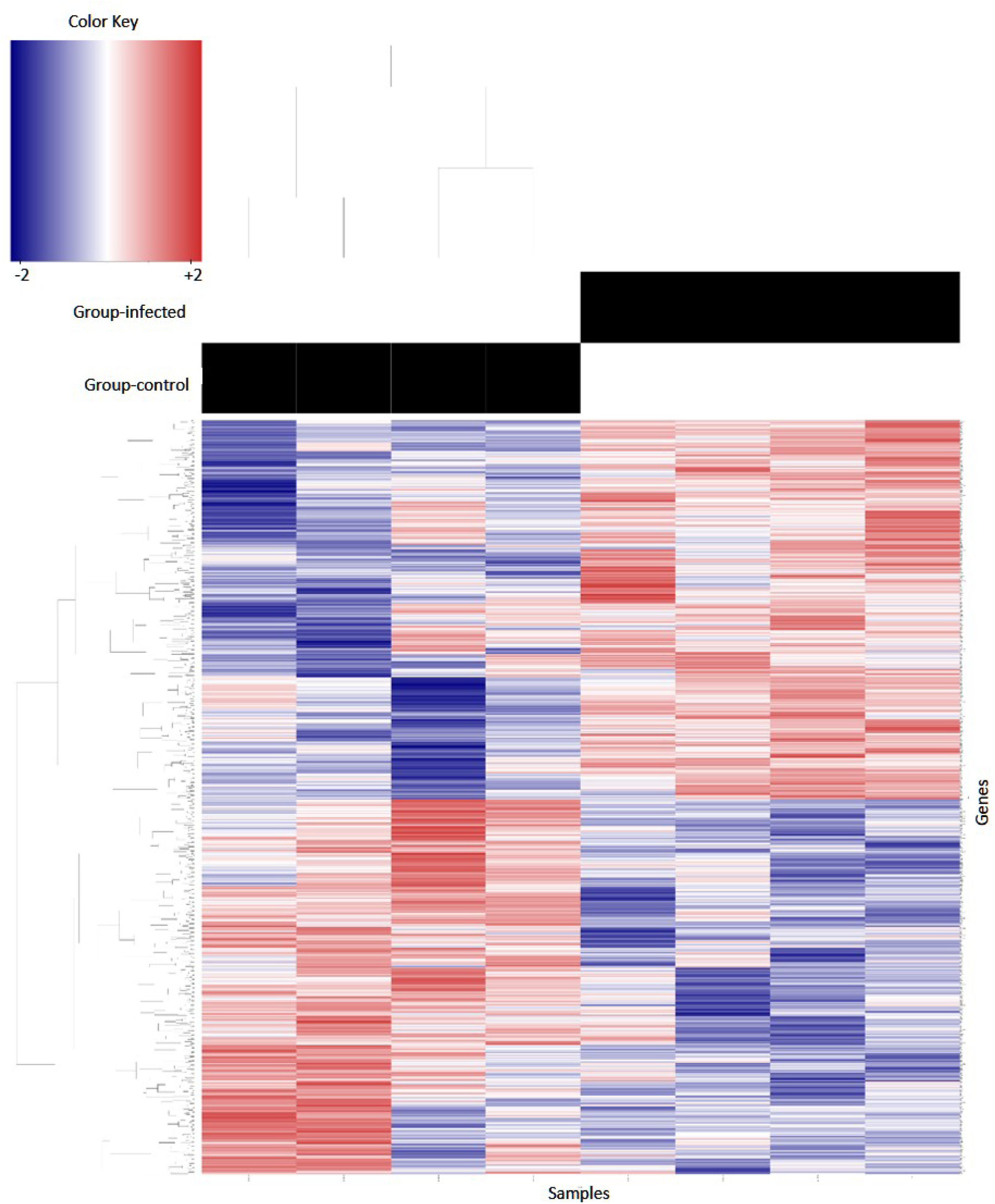
Expression heat map of up- and down-regulated mouse fetal heart genes at E14 comparing controls CVB3-infected and. All 13,118 protein-coding genes detected by RNA-Seq analysis were plotted using the color blue for down-regulation and red for up-regulation between and control (left 4 panels) CVB3-infected (right 4 panels) groups.

## REFERENCES

1. Triedman, J.K. and J.W. Newburger, Trends in congenital heart disease. Circulation, 2016. 133(25): p. 2716–2733.

2. Spector, L.G., J.S. Menk, J.H. Knight, C. McCracken, A.S. Thomas, J.M. Vinocur, M.E. Oster, J.D. St Louis, J.H. Moller, and L. Kochilas, Trends in Long-Term Mortality After Congenital Heart Surgery. Journal of the American College of Cardiology, 2018. 71(21): p. 2434–2446.

3. Triedman, J.K. and J.W. Newburger, Trends in Congenital Heart Disease: The Next Decade. Circulation, 2016. 133(25): p. 2716–33.

4. Vaughan, C.J. and C.T. Basson, Molecular determinants of atrial and ventricular septal defects and patent ductus arteriosus. American Journal of Medical Genetics - Seminars in Medical Genetics, 2000. 97(4): p. 304–309.

5. Yazigi, A., A.E. De Pecoulas, C. Vauloup-Fellous, L. Grangeot-Keros, J.M. Ayoubi, and O. Picone, Fetal and neonatal abnormalities due to congenital rubella syndrome: a review of literature. Journal of Maternal-Fetal and Neonatal Medicine, 2017. 30(3): p. 274–278.

6. Kleinman, D.S., B.D. Poole, G.M. Beckman, M.F. Hammersly, and T.A. Montgomery, Rubella. Some comments on the 1964-65 epidemic in California. California medicine, 1968. 109(4): p. 279–285.

7. Racicot, K. and G. Mor, Risks associated with viral infections during pregnancy. Journal of Clinical Investigation, 2017. 127(5): p. 1591–1599.

8. Stegmann, B.J. and J.C. Carey, TORCH Infections. Toxoplasmosis, Other (syphilis, varicella-zoster, parvovirus B19), Rubella, Cytomegalovirus (CMV), and Herpes infections. Current women”s health reports, 2002. 2(4): p. 253–258.

9. Liang, Q., W. Gong, D. Zheng, R. Zhong, Y. Wen, and X. Wang, The influence of maternal exposure history to virus and medicine during pregnancy on congenital heart defects of fetus. Environ Sci Pollut Res Int, 2017. 24(6): p. 5628–5632.

10. Liang, B., J.P. Guida, M.L. Costa Do Nascimento, and I.U. Mysorekar, Host and viral mechanisms of congenital Zika syndrome. Virulence, 2019. 10(1): p. 768–775.

11. Moore, M., M.H. Kaplan, J. McPhee, D.J. Bregman, and S.W. Klein, Epidemiologic, clinical, and laboratory features of coxsackie B1-B5 infections in the United States, 1970-79. Public Health Reports, 1984. 99(5): p. 515–522.

12. Coyne, C.B. and J.M. Bergelson, CAR: A virus receptor within the tight junction. Advanced Drug Delivery Reviews, 2005. 57(6): p. 869–882.

13. Koi, H., J. Zhang, A. Makrigiannakis, S. Getsios, C.D. MacCalman, G.S. Kopf, J.F. Strauss Iii, and S. Parry, Differential expression of the coxsackievirus and adenovirus receptor regulates adenovirus infection of the placenta. Biology of Reproduction, 2001. 64(3): p. 1001–1009.

14. Bendig, J.W.A., O.M. Franklin, A.K. Hebden, P.J. Backhouse, J.P. Clewley, A.P. Goldman, and N. Piggott, Coxsackievirus B3 sequences in the blood of a neonate with congenital myocarditis, plus serological evidence of maternal infection. Journal of Medical Virology, 2003. 70(4): p. 606–609.

15. Modlin, J.F. and C.S. Crumpacker, Coxsackievirus B infection in pregnant mice and transplacental infection of the fetus. Infection and Immunity, 1982. 37(1): p. 222–226.

16. Jaidane, H., A. Halouani, H. Jmii, F. Elmastour, M. Mokni, and M. Aouni, Coxsackievirus B4 vertical transmission in a murine model. Virol J, 2017. 14(1): p. 16.

17. Neumann, D.A., J.R. Lane, A. LaFond-Walker, G.S. Allen, S.M. Wulff, A. Herskowitz, and N.R. Rose, Heart-specific autoantibodies can be eluted from the hearts of Coxsackievirus B3-infected mice. Clin Exp Immunol, 1991. 86(3): p. 405–12.

18. Brown, G.C. and T.N. Evans, Serologic evidence of Coxsackievirus etiology of congenital heart disease. Journal of the American Medical Association, 1967. 199(3): p. 183–187.

19. Euscher, E., J. Davis, I. Holzman, and G.J. Nuovo, Coxsackie virus infection of the placenta associated with neurodevelopmental delays in the newborn. Obstetrics and Gynecology, 2001. 98(6): p. 1019–1026.

20. Lansdown, A.B.G., Coxsackievirus B3 infection in pregnancy and its influence on foetal heart development. British Journal of Experimental Pathology, 1977. 58(4): p. 378–385.

21. Sharma, V., L.S. Goessling, A.K. Brar, and P. Eghtesady, In Utero Infection with Coxsackievirus-B is Associated with Congenital Pulmonary Atresia. medRxiv, 2020: p. 2020.04.18.20070813.

22. Klein, S.L. and S. Huber, Sex differences in susceptibility to viral infection, in Sex Hormones and Immunity to Infection. 2010. p. 93–122.

23. Tennant, P.W., S.D. Samarasekera, T. Pless-Mulloli, and J. Rankin, Sex differences in the prevalence of congenital anomalies: A population-based study. Birth Defects Research Part A - Clinical and Molecular Teratology, 2011. 91(10): p. 894–901.

24. Hierholzer, J.C. and R.A. Killington, Virus isolation and quantitation. Virology Methods Manual, 1996: p. 25–46.

25. McFarlane, L., V. Truong, J.S. Palmer, and D. Wilhelm, Novel PCR assay for determining the genetic sex of mice. Sexual Development, 2013. 7(4): p. 207–211.

26. Lee, L.H., H. Yang, and G. Bigras, Current breast cancer proliferative markers correlate variably based on decoupled duration of cell cycle phases. Scientific Reports, 2014. 4.

27. Halonen, P., E. Rocha, J. Hierholzer, B. Holloway, T. Hyypia, P. Hurskainen, and M. Pallansch, Detection of enteroviruses and rhinoviruses in clinical specimens by PCR and liquid-phase hybridization. Journal of Clinical Microbiology, 1995. 33(3): p. 648–653.

28. Hwang, J.Y., K.M. Lee, Y.H. Kim, H.M. Shim, Y.K. Bae, J.H. Hwang, and H. Park, Pregnancy loss following coxsackievirus B3 infection in mice during early gestation due to high expression of Coxsackievirus-Adenovirus Receptor (CAR) in uterus and embryo. Experimental Animals, 2014. 63(1): p. 63–72.

29. Chung, A.C.K., X.R. Huang, L. Zhou, R. Heuchel, K.N. Lai, and H.Y. Lan, Disruption of the Smad7 gene promotes renal fibrosis and inflammation in unilateral ureteral obstruction (UUO) in mice. Nephrology Dialysis Transplantation, 2009. 24(5): p. 1443–1454.

30. Wang, Y., B. Wu, A.A. Chamberlain, W. Lui, P. Koirala, K. Susztak, D. Klein, V. Taylor, and B. Zhou, Endocardial to Myocardial Notch-Wnt-Bmp Axis Regulates Early Heart Valve Development. PLoS ONE, 2013. 8(4).

31. Tsukamoto, S., T. Mizuta, M. Fujimoto, S. Ohte, K. Osawa, A. Miyamoto, K. Yoneyama, E. Murata, A. Machiya, E. Jimi, S. Kokabu, and T. Katagiri, Smad9 is a new type of transcriptional regulator in bone morphogenetic protein signaling. Scientific Reports, 2014. 4.

32. Chen, J.W., B. Zhou, Q.C. Yu, S.J. Shin, K. Jiao, M.D. Schneider, H.S. Baldwin, and J.M. Bergelson, Cardiomyocyte-specific deletion of the coxsackievirus and adenovirus receptor results in hyperplasia of the embryonic left ventricle and abnormalities of sinuatrial valves. Circulation Research, 2006. 98(7): p. 923–930.

33. Chen, P., Y. Xie, E. Shen, G.G. Li, Y. Yu, C.B. Zhang, Y. Yang, Y. Zou, J. Ge, R. Chen, and H. Chen, Astragaloside IV attenuates myocardial fibrosis by inhibiting TGF-β1 signaling in coxsackievirus B3-induced cardiomyopathy. European Journal of Pharmacology, 2011. 658(2-3): p. 168–174.

34. Millan, F.A., F. Denhez, P. Kondaiah, and R.J. Akhurst, Embryonic gene expression patterns of TGF beta 1, beta 2 and beta 3 suggest different developmental functions in vivo. Development, 1991. 111(1): p. 131–43.

35. Verheugt, C.L., C.S.P.M. Uiterwaal, E.T. Van Der Velde, F.J. Meijboom, P.G. Pieper, H.W. Vliegen, A.P.J. Van Dijk, B.J. Bouma, D.E. Grobbee, and B.J.M. Mulder, Gender and outcome in adult congenital heart disease. Circulation, 2008. 118(1): p. 26–32.

36. Liang, Q., W. Gong, D. Zheng, R. Zhong, Y. Wen, and X. Wang, The influence of maternal exposure history to virus and medicine during pregnancy on congenital heart defects of fetus. Environmental Science and Pollution Research, 2017. 24(6): p. 5628–5632.

37. Watson, W.J., S. Awadallah, and M. Jo Jaqua, Intrauterine Infection With Coxsackievirus: Is It a Cause of Congenital Cardiac Malformations? Infectious Diseases in Obstetrics and Gynecology, 1995. 3(2): p. 79–81.

38. Soike, K., Coxsackie B-3 virus infection in the pregnant mouse. Journal of Infectious Diseases, 1967. 117(3): p. 203–208.

39. Tian, T., Y. Yang, L. Zhou, F. Luo, Y. Li, P. Fan, X. Dong, Y. Liu, J. Cui, and X. Zhou, Left Ventricular Non-Compaction: A Cardiomyopathy With Acceptable Prognosis in Children. Heart Lung and Circulation, 2018. 27(1): p. 28–32.

40. Zhang, W., H. Chen, X. Qu, C.P. Chang, and W. Shou, Molecular mechanism of ventricular trabeculation/compaction and the pathogenesis of the left ventricular noncompaction cardiomyopathy (LVNC). Am J Med Genet C Semin Med Genet, 2013. 163C(3): p. 144–56.

41. Miszalski-Jamka, K., J.L. Jefferies, W. Mazur, J. Glowacki, J. Hu, M. Lazar, R.A. Gibbs, J. Liczko, J. Klys, E. Venner, D.M. Muzny, J. Rycaj, J. Bialkowski, E. Kluczewska, Z. Kalarus, S. Jhangiani, H. Al-Khalidi, T. Kukulski, J.R. Lupski, W.J. Craigen, and M.N. Bainbridge, Novel Genetic Triggers and Genotype-Phenotype Correlations in Patients With Left Ventricular Noncompaction. Circ Cardiovasc Genet, 2017. 10(4).

42. Khan, R. and R. Sheppard, Fibrosis in heart disease: Understanding the role of transforming growth factor-β1 in cardiomyopathy, valvular disease and arrhythmia. Immunology, 2006. 118(1): p. 10–24.

43. Saxon, J.G., D.R. Baer, J.A. Barton, T. Hawkins, B. Wu, T.C. Trusk, S.E. Harris, B. Zhou, Y. Mishina, and Y. Sugi, BMP2 expression in the endocardial lineage is required for AV endocardial cushion maturation and remodeling. Developmental Biology, 2017. 430(1): p. 113–128.

44. Prall, O.W.J., M.K. Menon, M.J. Solloway, Y. Watanabe, S. Zaffran, F. Bajolle, C. Biben, J.J. McBride, B.R. Robertson, H. Chaulet, F.A. Stennard, N. Wise, D. Schaft, O. Wolstein, M.B. Furtado, H. Shiratori, K.R. Chien, H. Hamada, B.L. Black, Y. Saga, E.J. Robertson, M.E. Buckingham, and R.P. Harvey, An Nkx2-5/Bmp2/Smad1 Negative Feedback Loop Controls Heart Progenitor Specification and Proliferation. Cell, 2007. 128(5): p. 947–959.

45. Anderson, R.H., S. Webb, N.A. Brown, W. Lamers, and A. Moorman, Development of the heart: (2) Septation of the atriums and ventricles. Heart, 2003. 89(8): p. 949–958.

46. Heallen, T., M. Zhang, J. Wang, M. Bonilla-Claudio, E. Klysik, R.L. Johnson, and J.F. Martin, Hippo pathway inhibits Wnt signaling to restrain cardiomyocyte proliferation and heart size. Science, 2011. 332(6028): p. 458–61.

47. Lyden, D., J. Olsewski, and S.A. Huber, Influence of sex hormones on coxsackie virus group B, type 3 induced myocarditis in Balb/c mice. European Heart Journal, 1987. 8(SUPPL. J): p. 389–391.

48. Coronado, M.J., J.E. Brandt, E. Kim, A. Bucek, D. Bedja, E.D. Abston, J. Shin, K.L. Gabrielson, W. Mitzner, and D. Fairweather, Testosterone and interleukin-1β increase cardiac remodeling during coxsackievirus B3 myocarditis via serpin A 3n. American journal of physiology. Heart and circulatory physiology, 2012. 302(8): p. H1726–H1736.

49. Miller-Hance, W.C. and T.A. Tacy, Gender differences in pediatric cardiac surgery: The cardiologist’s perspective. Journal of Thoracic and Cardiovascular Surgery, 2004. 128(1): p. 7–10.

50. Engelfriet, P. and B.J.M. Mulder, Gender differences in adult congenital heart disease. Netherlands Heart Journal, 2009. 17(11): p. 414–417.

51. Hadar, E., N. Melamed, M. Sharon-Weiner, S. Hazan, D. Rabinerson, M. Glezerman, and Y. Yogev, The association between stillbirth and fetal gender. Journal of Maternal-Fetal and Neonatal Medicine, 2012. 25(2): p. 158–161.

